# Improved Ethanol Tolerance and Production in Pyruvate Dehydrogenase Mutant of *Saccharomyces cerevisiae*

**DOI:** 10.1101/2023.01.29.526136

**Authors:** Anna Yang, Tahina O. Ranaivoarisoa, Arpita Bose

## Abstract

Ethanol, a naturally synthesized compound by *Saccharomyces cerevisiae* yeast through alcoholic fermentation, has previously been studied as a renewable alternative to traditional fossil fuels. However, current challenges of engineering *S. cerevisiae* strains for ethanol production remain: low ethanol productivity, inefficient substrate catabolism, and a buildup of toxic products to inhibitory levels. In this study, we proposed a method of metabolic rewiring via the deletion of the *pda1* gene, which leads to pyruvate dehydrogenase (PDH) deficiency. The Δ*pda1* mutant strain was created by CRISPR Cas-9 knockout using the constructed pCRCT-PDA1 plasmid. Subsequently, mutant candidates were screened by PCR and Sanger sequencing, confirming a 17 bp deletion in the *pda1* gene. The wild-type and mutant strains were analyzed for growth under aerobic and anaerobic conditions in glucose and glycerol, as well as ethanol production and tolerance. The Δ*pda1* mutant displays a ~two-fold increase in anaerobic ethanol production and an aerobic growth defect with no observed increase in ethanol production. The mutant is also hyper-tolerant to ethanol, which allows a faster buildup of products in growth media with minimal reduction in growth. This new *S. cerevisiae* strain deficient in PDH may provide a solution to the efficient and abundant synthesis of biofuels such as ethanol by redirecting metabolic flux and altering stress response.

## Introduction

The consumption of fossil fuels and increasing greenhouse gas emissions that accompanies fuel combustion demand the developments of more sustainable alternatives.^1^ Ethanol is a carbon-neutral, renewable biofuel that has been studied as an alternative to traditional fossil fuels to decrease carbon emissions.^1^ Despite its numerous environmental benefits, further efficient and affordable synthesis of ethanol remains an obstacle to more competitive commercialization.^2^ *Saccharomyces cerevisiae* yeast is characterized by its split aerobic/anaerobic metabolic pathways, specifically its ability to produce ethanol through alcoholic fermentation in the presence of a carbon source such as glucose, glycerol, or cellulose — glucose being the preferred carbon source for fermentation.^3^ Previous literature suggests that even under fully aerobic conditions, glucose excess causes yeast to repress its aerobic respiration pathway in favor of fermentation in a process known as the Crabtree effect.^4^ This is due to an energetic overflow in sugar metabolism, an evolutionarily favorable process to accelerate glucose intake under limited respiratory capacities. However, the efficiency of glucose to ethanol conversion in wild-type yeast is suboptimal and is further hindered by the toxicity of ethanol to microbes including yeast. The small, amphipathic nature of ethanol molecules enhances its permeability through cell membranes. High concentrations of ethanol in a growth medium hinder cellular growth and division by disturbing protein folding and thus the metabolic activity of yeast cells.^5^

Modifying metabolic pathways to increase ethanol production rate has been studied extensively, but previous studies have focused primarily on downregulating genes involved in glycerol synthesis (GPD, GDH, ADH, and PDC) to improve ethanol yield at the expense of biomass production.^6^ However, glycerol and other biomass compounds are crucial for cellular processes such as osmoregulation, and their hindrance results in decreased growth rates in mutant strains.^6^

Pyruvate is a key intermediate located at the entrance of the citric acid cycle, which acts as a gateway between glycolysis and aerobic respiration. The pyruvate dehydrogenase complex (PDH), its E1α subunit encoded by the *pda1* gene, catalyzes the decarboxylation of pyruvate into acetyl-CoA, which is a precursor to oxidation in the aerobic respiration pathway. In the case of a PDH mutation, the combined expression of *PDC, ADH*, and *ACS* genes allows for limited acetyl-CoA synthesis via the PDH bypass.^7^ However, the relatively low affinity of cytosolic pyruvate decarboxylase (PDC) for pyruvate compared to pyruvate dehydrogenase (PDH) indicates a preference for pyruvate metabolism through the PDH complex.^8^ We observed that the deletion of the *pda1* gene disrupts PDH function, directing glucose intake toward alcoholic fermentation without completely abolishing aerobic respiration, the primary source of ATP production in respiratory yeast cells. By identifying this metabolic-synthetic balance in PDH-negative *S. cerevisiae*, we were able to increase ethanol yield while maintaining a stable specific growth rate under anaerobic conditions.

## Methods

### Strain, Media, and Growth Conditions

The *S. cerevisiae* strain BY4741 (MATa his3Δ1 leu2Δ0 met15Δ0 ura3Δ0) is an S288C-derivative laboratory strain used in this study.^9^ Yeast cultures were grown in a standard YPD medium (1% yeast extract, 2% tryptone, 0.5% glucose) at 30°C with a shaking speed of 200 rpm pre-transformation. After transformation with the uracil (URA) containing plasmid pCRCT-*PDA1*, cells were selected for plasmid incorporation by growing for 48 hours on a synthetic complete dropout uracil (SC-URA) agar plate. Pre-cultures were grown overnight in 5 mL of YPD medium with 0.5% glucose before each growth experiment.

### Construction of the Recombinant pCRCT-PDA1 Plasmid

The pCRCT plasmid was used to perform single gene homology-integrated CRISPR-Cas9 knockout in *S. cerevisiae* BY4741 as described in the methods of Bao.^10^ The pCRCT plasmid was a gift from Huimin Zhao (Addgene plasmid # 60621; http://n2t.net/addgene:60621; RRID: Addgene_60621) and purchased from Sigma.

The donor insert fragments (132 bp, 142 bp, 132 bp) were constructed by two rounds of overlap extension PCR (OE-PCR) with the listed primers (Table 1). The first round of thermocycling constructs the template from short fragments with overlap regions. In the second round of thermocycling, the template was amplified with primers F1 and R3 for a second round of PCR reactions to amplify the donor fragments.

**Table 1.**
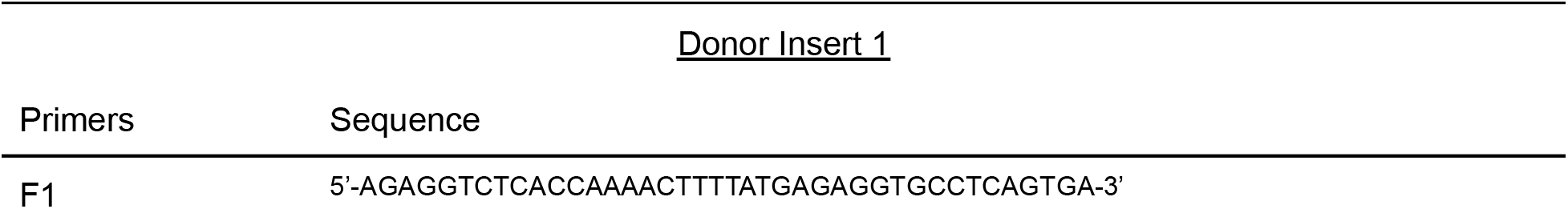
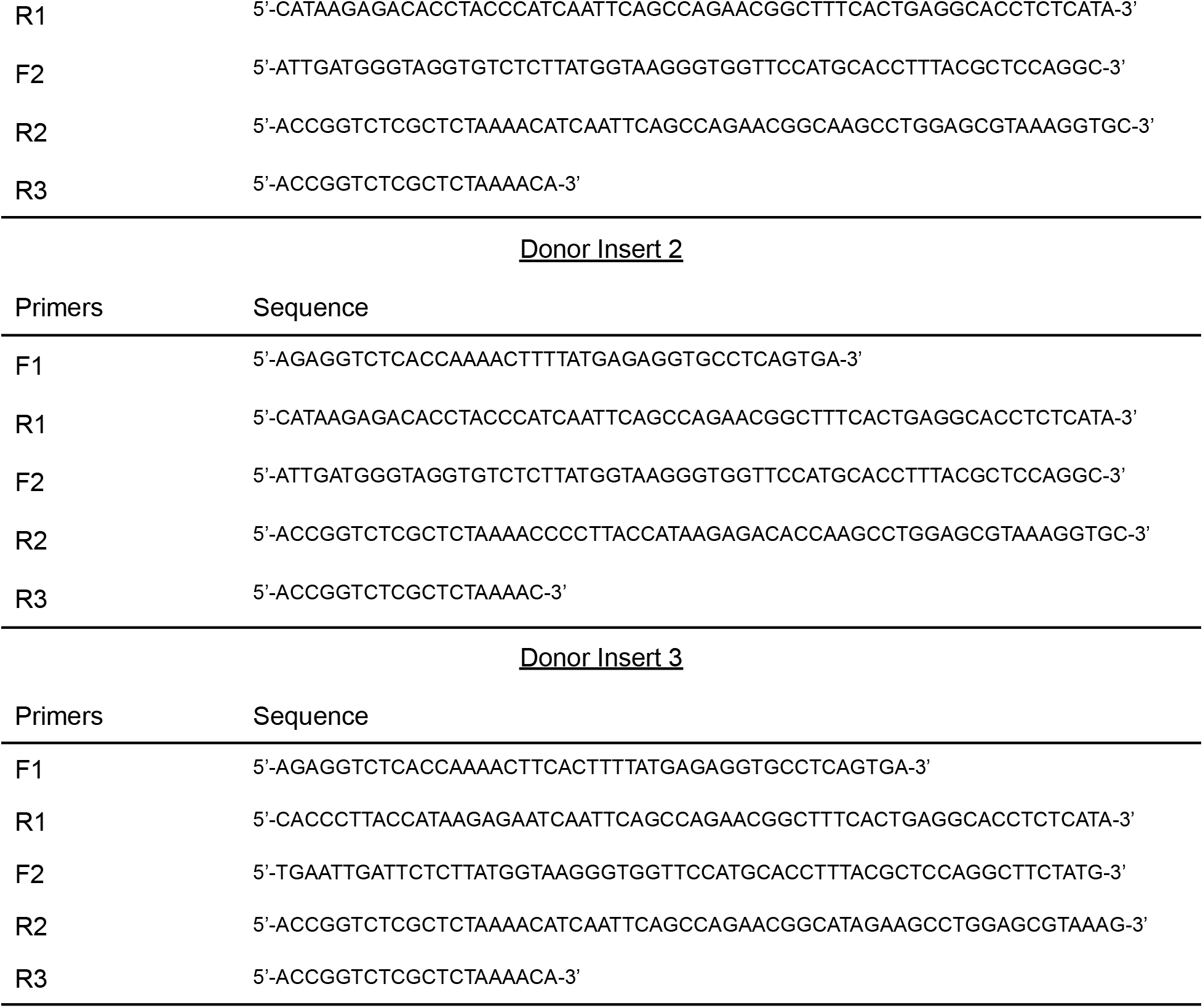
Primers used to construct the candidate *pda1* donor inserts by OE-PCR.

The resulting insert DNA is cloned into the pCRCT plasmid through Golden Gate Assembly. The pCRCT plasmid was digested with the *Bsa*I restriction enzyme at two sites to remove a 459 bp fragment. Donor insert fragments 1, 2, and 3 were independently cloned into three *Bsa*I digested pCRCT samples and ligated with T4 ligase to assemble the recombinant plasmids (Fig. 1A). The pCRCT-*PDA1* plasmids were amplified by transformation into *E. coli* DH5α cells made competent through calcium chloride washes. The competent cells were heat shocked at 42°C for 30 seconds and recovered on ice for 5 minutes. The transformed cell suspension was plated on an ampicillin LB medium for selection. Colony PCR and sequencing were performed to verify successful transformants (Fig. 1C).

**Figure 1.**
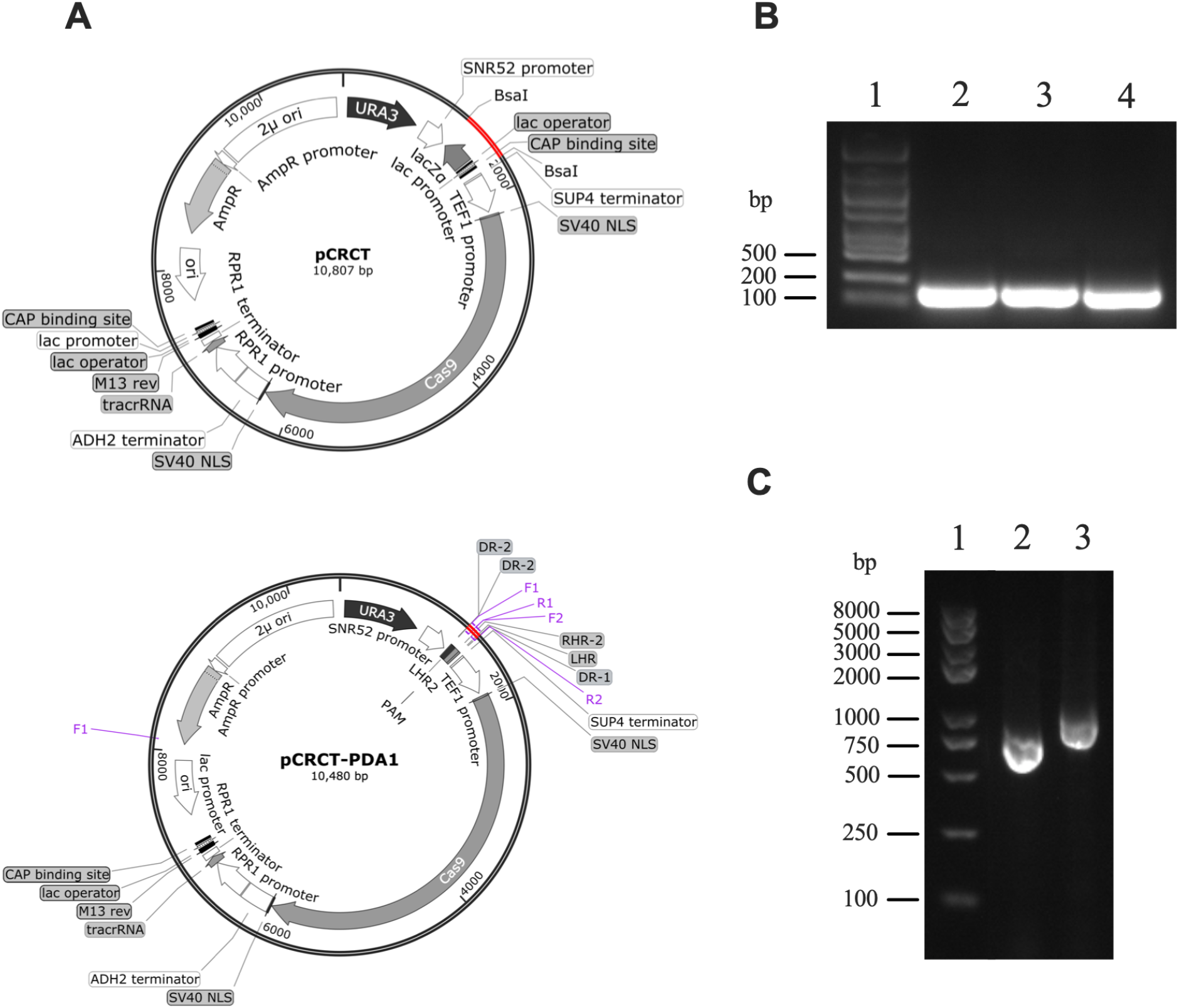
Construction and replication of the recombinant pCRCT-*PDA1* plasmid. **(A)** Plasmid maps of the pCRCT and pCRCT-*PDA1* plasmids. Red regions represent the replaced and inserted fragments. **(B)** Gel electrophoresis of 3 candidate donor fragments constructed by overlap extension PCR (OE-PCR). Lane 1, RB-MK8 DNA marker; Lanes 2-4, *pda1* donor insert fragments 1, 2, and 3. **(C)** Gel electrophoresis following colony PCR using pCRCT VF and VR primers in post-transformation *E. coli* DH5α. Lane 1, RB-MK8 DNA marker; Lane 2, 750 bp sequence of pCRCT-PDA1 (DI 3) in successful transformants; Lane 3, cells transformed with the original pCRCT plasmid (positive control).

Transformed *E. coli* cells were grown in a liquid LB medium with 0.1% ampicillin overnight at 37°C. The plasmid extraction protocol was performed on the overnight-incubated cell cultures, resulting in pCRCT-*PDA1* plasmid stocks that can be used or stored at −80°C.

### Disruption of the pda1 Gene in BY4741 Yeast by CRISPR-Cas9

pCRCT-*PDA1* plasmids were transformed into *S. cerevisiae* BY4741 using the LiAc/SS carrier DNA/PEG method developed by Gietz and Schiestl.^11^ Three serial dilutions of the transformed cell suspension (200 μl, 20 μl, and 2 μl) were plated on synthetic complete dropout uracil (SC-URA) agar plates for selection. A negative control with wild-type BY4741 yeast and a positive control with cells transformed with pCRCT were also plated to ensure that the transformation was successful and an absence of colonies for Δ*pda1* yeast can be attributed to the mutation. Colonies were selected for colony PCR amplification of the *pda1* gene and sequenced for a 17 bp deletion that introduces a premature stop codon (p.Met182IlefsTer186). The mutant was verified for successful disruption using PCR and Sanger sequencing of PDH by sequence alignment.

### Growth Assays

To determine the effect of carbon source concentration on the specific growth rates of PDH-negative *S. cerevisiae*, cells were inoculated in 200 μL of YPD medium with glucose (0.1%, 0.5%, 1%, 2.5%, 5%, 10%, and 20%) or glycerol (0.5%) as their sole carbon source. Ethanol was added to 0.5% glucose cultures in various concentrations (1%, 2%, 5%, 8%, and 10%) to measure ethanol stress response and tolerance in Δ*pda1* and wild-type cells under a standardized carbon supply. The cells were cultured in 96-well microplates at 24°C with a shaking speed of 200 rpm and optical density measurements were taken every 15 minutes at 600 nm over a 48-hour incubation period. Growth rates were calculated by averaging the maximum specific growth rates (μ_max_) of five independently cultured biological replicates during exponential growth (*n*=5). All assays were performed aerobically and then repeated under anaerobic conditions achieved by purging the medium in anaerobic culture vials with anoxic N_2_/CO_2_ for 5 minutes and vacuuming for 10 minutes. All anaerobic inoculations were performed in an anaerobic chamber and the 96-well plates were sealed using microplate sealing film throughout growth to ensure the absence of oxygen.

### Ethanol Colorimetric Assays

Aerobic and anaerobic ethanol production of Δ*pda1* and wild-type cultures were measured for glucose concentrations of 0.5%, 2.5%, 5%, 10%, and 20% using the Ethanol Assay Kit provided by BioAssay Systems (catalog no. ECET100; BioAssay Systems, Hayward, CA, USA). 3.33 μL of cell cultures with an initial OD of 1.8 were inoculated into 200 μL of liquid YPD medium in each well of a 96-well microplate for 48 hours at 24°C, 200 rpm. After 48 hours of growth, the supernatant was extracted and diluted with ethanol assay buffer to make a 10% solution, and enzymatic assays were performed according to the Ethanol Assay Kit protocol from BioAssay Systems. Ethanol concentrations were estimated by measuring absorbance at 670 nm after incubating at 37°C for 30 minutes and converted to nmol/μL using a standard curve. Concentrations were an average of 5 replicates graphed as percentages (*n*=5).

### Statistical Analysis

Statistical analysis was conducted on the growth and ethanol production data to determine significance. Unpaired, two-tailed t-tests were performed between mutant and wild-type μ_max_ averages for all concentrations of glucose, glycerol, or ethanol to confirm growth differences. Similarly, t-tests were performed between mutant and wild-type ethanol production averages to indicate a significant production increase in mutant cells. This statistical method was chosen because of non-categorical data and independent groups. The cut-off *p*-value for significance is defined as *p*=0.05, which indicates a meaningful difference between the two experimental groups.

## Results and Discussion

### Growth Defects in Δpda1 Mutant Under Aerobic Conditions

To examine the aerobic and anaerobic growth differences between the wild-type and Δ*pda1* mutant strains, the specific growth rates were calculated using growth curves graphed from OD_600_ measurements. Figure 2A shows growth curves of mutant and wild-type cells grown aerobically under 1% glucose, to specifically emphasize the impact of the *pda1* deletion. Grown aerobically under 1% glucose, the mutant initially demonstrated slower growth during exponential phase but eventually plateaued at a higher OD than the wild type (*p*<0.01) (Fig. 2A). After 48 hours, the mutant reached a final OD of 1.68, while the wild type only reached an OD of 1.40 (Fig. 2A). While a reduced specific growth rate implies a deficiency in cellular metabolic rate, a higher maximum OD suggests a potential increase in toxicity tolerance because cell division was impeded less by a buildup of ethanol. A broader range of glucose concentrations (0.1% - 20%) was represented in Figures 2B and 2C. Under aerobic conditions, the mutant displayed growth defects with decreased specific growth rates for all glucose concentrations examined (*p*<0.001) (Fig. 2B). The results suggest that a deficiency in the pyruvate dehydrogenase enzyme causes low acetyl-CoA concentrations and ATP synthesis in the cell, decreasing aerobic metabolism in the mutant strain. Anaerobically, the growth difference between the two strains is insignificant (*p*>0.05) (Fig. 2C). The mutant displays a significant growth advantage in anaerobic conditions for all glucose concentrations examined (*p*<0.001), while the wild type shows no preference for either environment for glucose concentrations under 5% (*p*>0.05), and only shows anaerobic growth advantages for glucose concentrations over 5% (*p*<0.001) (Fig. 2). Both the wild type and mutant strains’ growth rates peak at 5% glucose concentration regardless of the presence of oxygen, after which the rate plateaus, and decreases due to a limit in the rate at which glucose can be metabolized.

**Figure 2.**
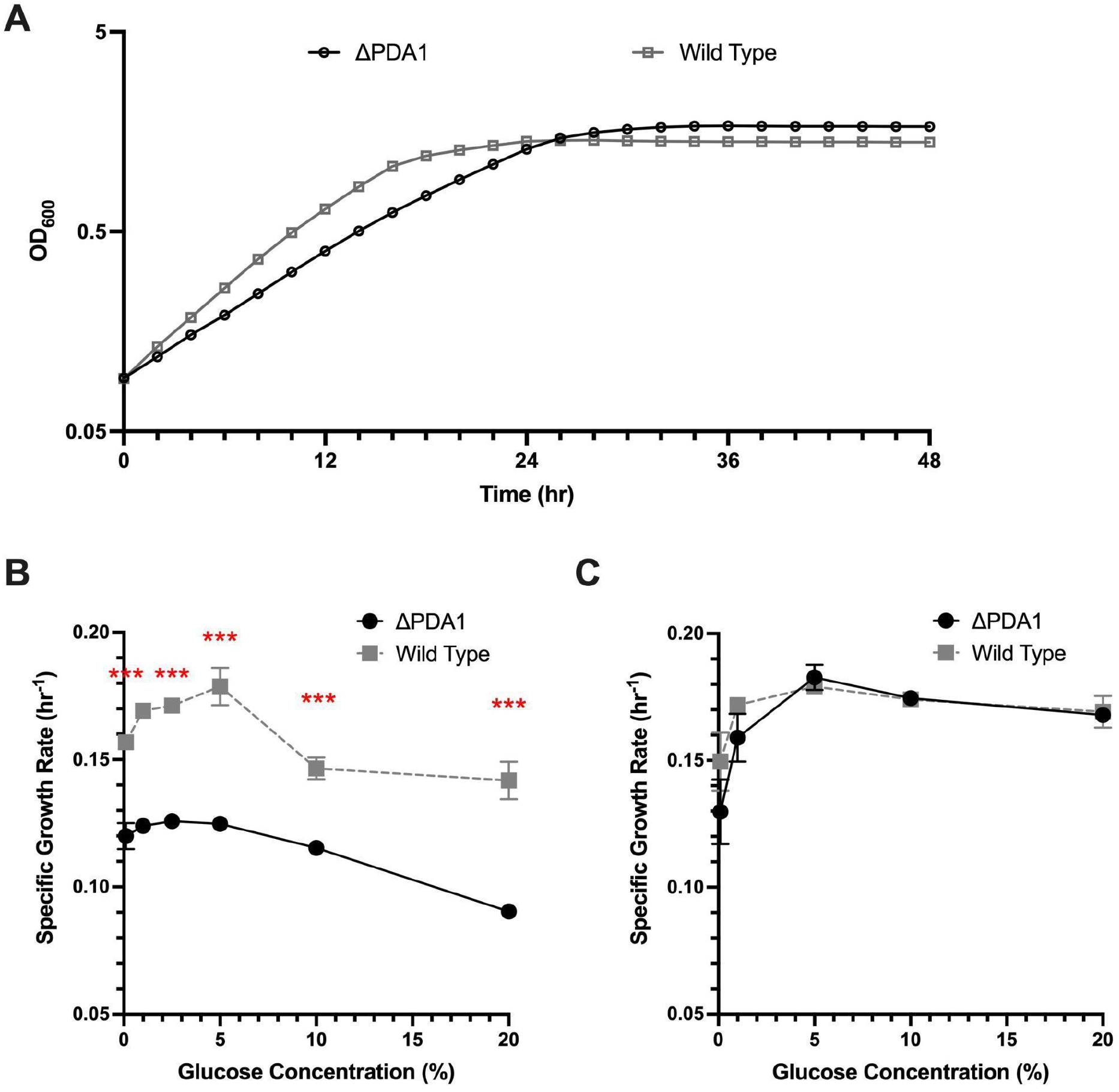
Specific growth rates of yeast cultures under various glucose concentrations and oxygen availability. **(A)** Growth curves graphed as OD_600_ values taken over 24 hours for the Δ*pda1* mutant strain in comparison to the wild type BY4741 yeast grown aerobically in YPD containing 1% glucose (*n*=5). **(B and C)** Specific growth rates (hr^−1^) of the mutant and wild type strains grown with 0.1% - 20% glucose under aerobic (B) and anaerobic (C) conditions. Specific Growth rates (hr^−1^) were calculated using OD_600_ measurements and averaged across 5 biological replicates (*n*=5). Uncertainty bars represent ±SD. Significance is determined by unpaired, two-tailed t-tests (\**p*<0.05, \*\**p*<0.01, \*\*\**p*<0.001).

One source of uncertainty during growth is the oxygen removal process for the anaerobic cultures. Anaerobic media was created and inoculated in the anaerobic chamber, but the growth process in the microplate reader may have let small amounts of oxygen into the sealed wells of a microplate. This could be responsible for the marginal differences between the two strains observed during anaerobic growth. Another source of uncertainty is that differential ethanol production was not considered when measuring growth rates. Due to a higher ethanol concentration present in the mutant cultures (see the following section), the ethanol in the medium could not be controlled, thus introducing a glucose-independent factor negatively affecting mutant growth rates.

### Mutants Lacking PDH Display Increased Ethanol Production in Anaerobic and High-Glucose Aerobic Conditions

The effect of the *pda1* gene deletion on yeast metabolism was determined by comparing the net ethanol production of mutant and wild-type cells with a loss of function mutation. Ethanol production was quantified colorimetrically in cell culture media after 24 hours of growth in both aerobic and anaerobic conditions (Fig. 3). Aerobically, the mutant demonstrated a marginally higher ethanol yield only when the glucose concentration was high; for example, under 10% glucose (*p*=0.023). When grown at lower glucose concentrations, there was no significant difference in ethanol production between the strains (*p*>0.05) (Fig. 3A). Anaerobically, however, the mutant consistently produced more ethanol regardless of the glucose concentration in the medium (*p*<0.05) (Fig. 3B). Anaerobic ethanol production in Δ*pda1* mutants demonstrated an increase ranging from 45.1% to 100.7% compared to the wild type yeast for all glucose concentrations examined (Fig. 3B). Anaerobic ethanol production in wild type cells plateaus at 10%, as there is no difference in production between 10% and 20% glucose (*p*>0.05), whereas Δ*pda1* mutants grown with 20% glucose still showed a 16.97% increase in ethanol yield compared to cultures grown at 10% glucose (*p*<0.05). This delayed plateau in mutant fermentation suggests a heightened capacity for Δ*pda1* cells to metabolize glucose anaerobically.

**Figure 3.**
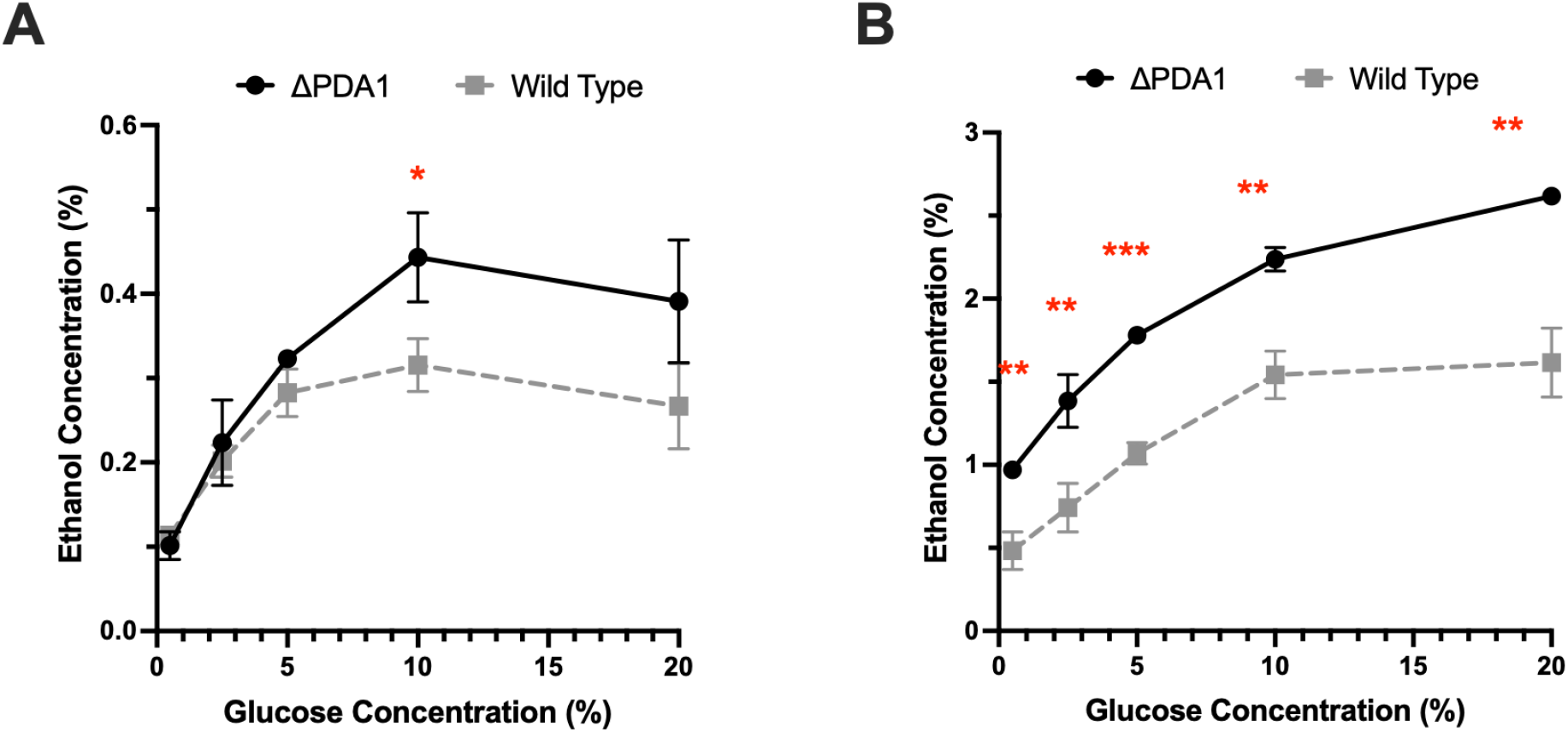
Ethanol production of mutant and wild-type yeast cultures fermented under various glucose concentrations. Ethanol concentration in solution was assayed colorimetrically by OD_670_ measurements after performing enzymatic reactions on supernatants from aerobic cultures (A) and anaerobic cultures (B) after 48 hours of growth in media containing 0.5% to 20% glucose. Ethanol production was averaged across 5 biological replicates (*n*=5). Uncertainty bars represent ±SD. Significance is determined by unpaired, two-tailed t-tests (\**p*<0.05, \*\**p*<0.01, \*\*\**p*<0.001).

The observed increase in ethanol production in Δ*pda1* mutants is perhaps due to a redirection of metabolic flux in the absence of the pyruvate dehydrogenase (PDH) enzyme. Located at the entrance of the citric acid cycle, PDH catalyzes the decarboxylation of pyruvate into acetyl-CoA, which acts as a gateway to the aerobic breakdown of glucose through cellular respiration (Fig. 4). A loss of function mutation in the E1α subunit, encoded by the *pda1* gene, causes the PDH enzyme to become dysfunctional and unable to convert pyruvate to acetyl-CoA. The buildup of excess pyruvate in the cell is metabolized anaerobically into ethanol. Thus, ethanol production in the mutant is higher both in anaerobic conditions, and aerobic conditions under an excess supply of glucose (Fig. 3). To compensate for the lack of PDH function, a PDH bypass mechanism is able to synthesize a limited amount of acetyl-CoA in Δ*pda1* mutants.^7^ This reaction, catalyzed by the *PDC, ADH*, and *ACS* genes, converts pyruvate to acetaldehyde, to acetate, and finally, to acetyl-CoA. The final acetate to acetyl-CoA reaction requires the hydrolysis of ATP, unlike the direct decarboxylation of pyruvate into acetyl-CoA by PDH.^7^ Because Δ*pda1* mutants energetically prefer the PDH pathway over the bypass, the strain is fully respiratory only when glucose supply is limited (Fig. 3A).

**Figure 4.**
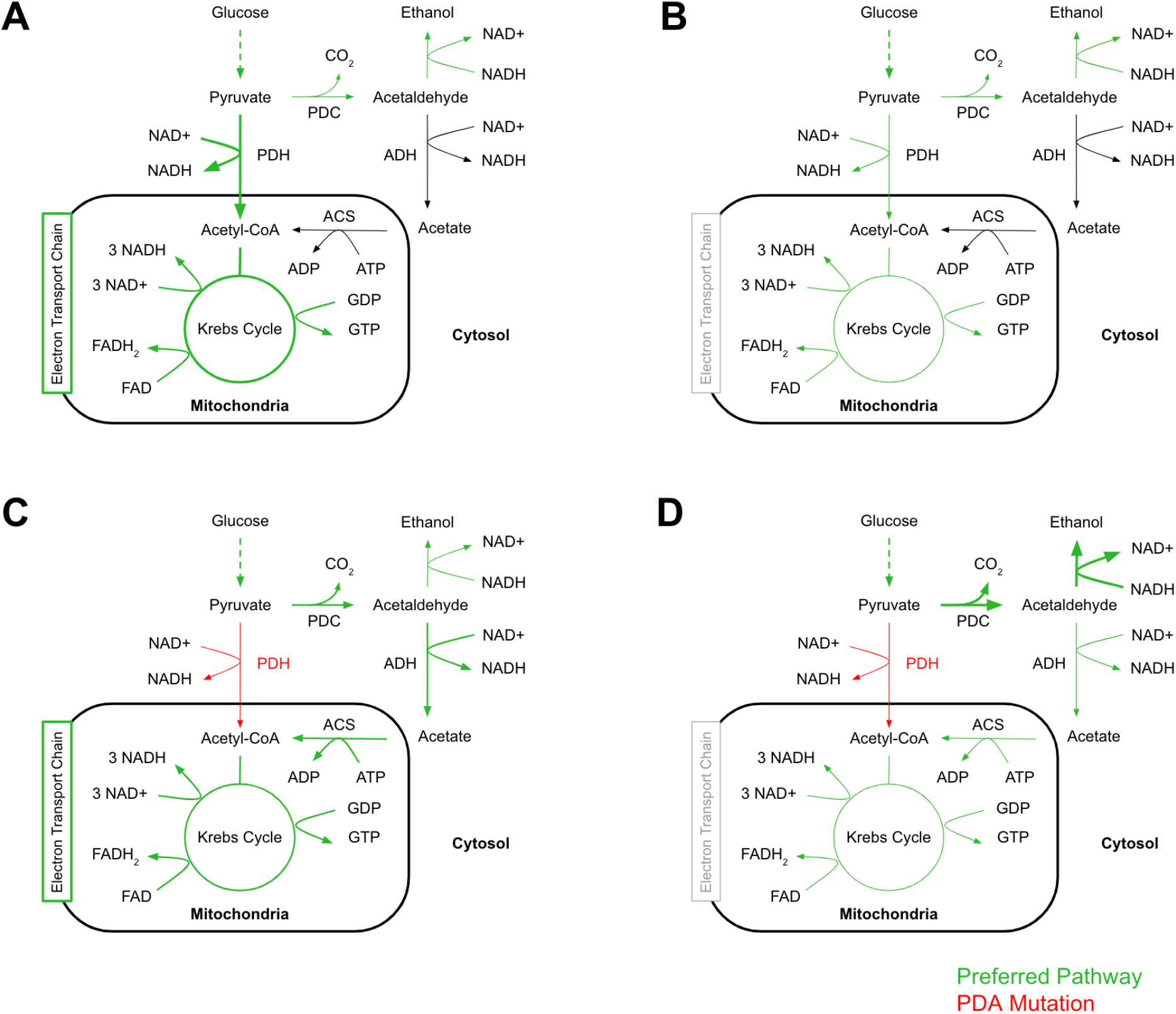
Proposed pathways of glucose metabolism in wild-type and Δ*pda1* mutant yeast. Metabolic flow was analyzed for cells under wild-type aerobic (A), wild-type anaerobic (B), mutant aerobic (C), and mutant anaerobic (D) growth conditions. The conversion of pyruvate to acetyl-CoA is completed either through PDH or the PDH bypass. Preferred pathways are denoted in green, and the product conversion rate is further distinguished by thickness. Red arrows denote a loss of function mutation of the PDH enzyme in Δ*pda1* mutants.^12,13^

During aerobic growth, wild-type cells prefer to metabolize glucose via the aerobic respiration pathway through the Krebs cycle and electron transport chain. The conversion of pyruvate to acetyl-CoA, a crucial step at the entrance of the Krebs cycle, can be completed either through PDH or the PDH bypass, which is a combined pathway utilizing the pyruvate decarboxylase (PDC), aldehyde dehydrogenase (ADH), and acetyl-CoA synthetase (ACS) genes (Fig. 4). Wild type cells use both functional pathways aerobically but prefer the direct conversion through PDH (Fig. 4A). The mutant is forced to use the bypass under aerobic conditions to continue using the electron transport chain, which is preferred to fermentation due to its high ATP output (Fig. 4C). However, because the bypass operates slower, ATP synthesis is impeded, reducing mutant growth rate. There are marginal differences in aerobic ethanol production, but mostly insignificant because fermentation doesn’t generate the ATP needed for the cells to grow.

During anaerobic growth, however, the electron transport chain is dysfunctional lacking oxygen as the ultimate electron acceptor. Therefore, the combined pathways of glycolysis and Krebs cycle contribute to the buildup of NADH and the lack of NAD^+^, which needs to be regenerated through fermentation to maintain redox balance. An increase in ethanol production allows the more efficient conversion of NADH to NAD^+^. In Δ*pda1* mutants, this process is preferred to the energetically unfavorable PDH bypass and allows glycolysis to continue by reducing the buildup of cellular NADH.

Comparing the ethanol production and growth rates of the Δ*pda1* mutant yeast provides insight into the optimal growth conditions for fermentation. Culturing Δ*pda1* mutant cells anaerobically at 5% glucose experimentally produces 69.66% of the theoretical yield, whereas the wild type produces only 41.82% of the theoretical yield (Fig. 3B). At 5% glucose, this 66.55% increase in experimental ethanol production for Δ*pda1* mutants accompanies an optimized anaerobic growth rate of 0.183 hr^−1^ (Fig. 2A).

The major source of uncertainty in analyzing the cause of the ethanol production increase is the lack of differentiation between the mutant pathway redirection (Fig. 4) and the possibility of increased ethanol tolerance in mutants (Fig. 5). The proposed pathways do not clearly indicate whether the observed difference in production is mainly due to an intrinsic change in metabolic flux favoring fermentation, or if the higher mutant OD reached in anaerobic conditions contributes to the ethanol production. Further studies could be conducted to pinpoint the specific cause of the excess ethanol produced. Mapping protein expression could confirm the proposed pathway changes in Figure 4 or point to a change in expression in another gene that contributes to ethanol tolerance.

**Figure 5.**
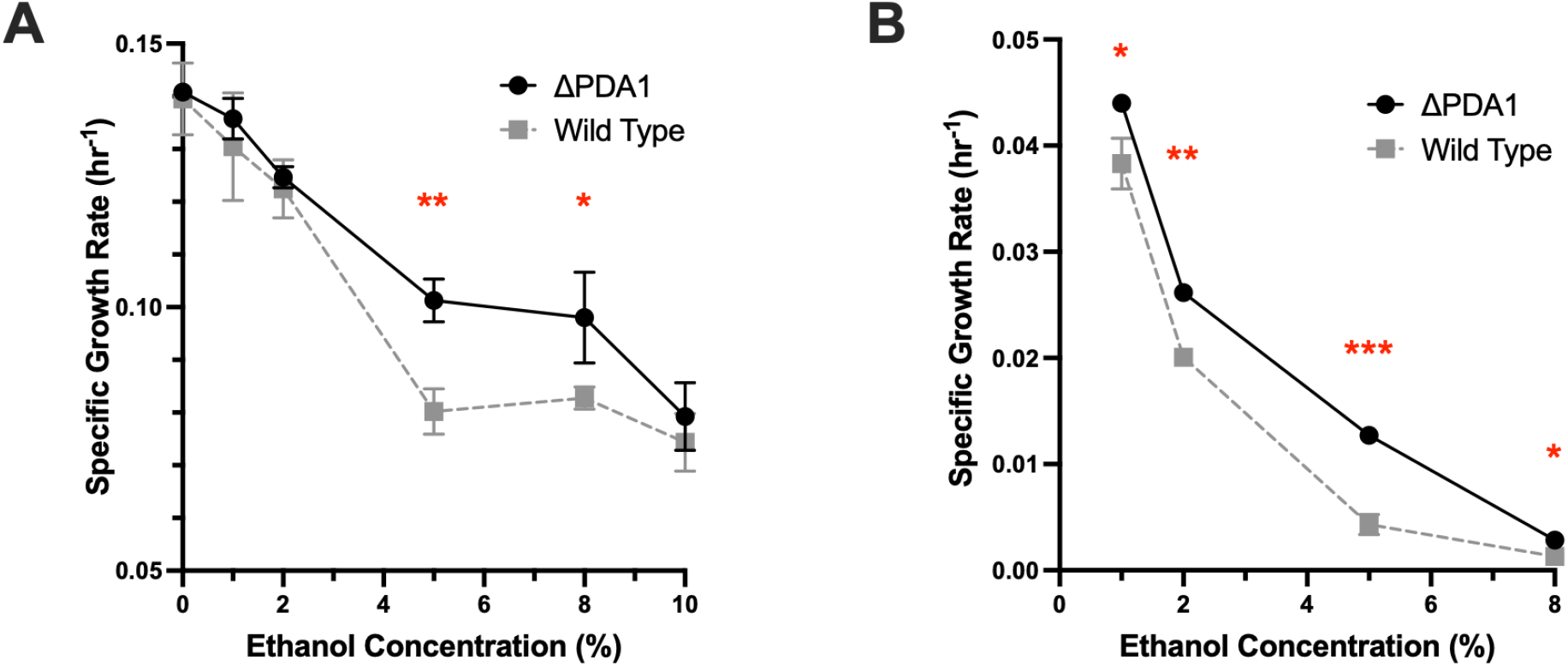
Ethanol tolerance of yeast cultures exposed to various concentrations of ethanol, measured by specific growth rates. Specific growth rates (hr^−1^) of the Δ*pda1* mutant strain in comparison to the wild-type BY4741 yeast grown in YPD containing 1% - 10% ethanol under aerobic (A) and anaerobic (B) conditions. Growth rates were calculated using OD_600_ measurements and averaged across 5 biological replicates (*n*=5). Uncertainty bars represent ±SD. Significance is determined by unpaired, two-tailed t-tests (\**p*<0.05, \*\**p*<0.01, \*\*\**p*<0.001).

### Increased Ethanol Tolerance in Δpda1 Mutants

Tolerance to various concentrations of ethanol in growth media is measured through OD_600_ measurements, from which the maximum specific growth rate is calculated as a proxy for ethanol resistance. Aerobically, Δ*pda1* mutants displayed a significant growth advantage in media containing around 5% - 8% ethanol (Fig. 4A). Lower and higher concentrations yielded no significant difference in growth rate (*p*>0.05). Anaerobically, Δ*pda1* mutants were more ethanol resistant compared to the wild type in all ethanol-containing environments observed (*p*<0.05) (Fig. 4B). Both strains generally demonstrated reduced growth as concentration increases, due to the lethality of ethanol against yeast cells. At concentrations at or exceeding 10% ethanol, cells were unable to grow anaerobically (Fig. 4B). Glucose abundance was controlled at 0.5% in all cultures to ensure growth rate remained independent of glucose availability.

Ethanol tolerance, especially during anaerobic growth, is important for increasing net ethanol production. Δ*pda1* mutants are able to reach a higher OD compared to wild-type yeast in sublethal ethanol-containing environments, and the increased cell viability allows an overall increase in ethanol synthesis rate. According to the data, the growth advantage of Δ*pda1* mutants was not counteracted by the increase in ethanol concentration in growth media over time resulting from a higher fermentation rate, as mutant cells continue to grow faster even as more ethanol accumulates in the medium (Fig. 3 and 5). The increase in ethanol tolerance is hypothesized to be attributed to the intensified selective pressure among Δ*pda1* mutants favoring ethanol-resistant cells. Due to a higher fermentation rate and thus, higher net production of ethanol, only tolerant Δ*pda1* cells were able to survive and pass on their acquired resistance to future generations. Another possibility is that the Δ*pda1* mutant upregulates alcohol dehydrogenase (ADH) expression due to an increased need to convert acetaldehyde to acetate as a step of the PDH bypass, reducing the buildup of the toxic aldehyde in the cell.

### Growth Advantage of Δpda1 Mutants Grown Aerobically in Glycerol

When grown aerobically with 0.5% glycerol as the sole carbon source, *Δpda1* mutants show a significant growth advantage over the wild type (*p*=0.0083), but this difference is not observed when both strains are grown in 0.5% glucose (*p*>0.05) (Fig. 6A). Both strains preferred metabolizing glucose over glycerol, however, demonstrated by their higher specific growth rates for glucose (Fig. 6A). Neither strains showed growth when grown with glycerol anaerobically, as opposed to the fermentable sugar glucose (Fig. 6B). When exposed to 5% ethanol in addition to the standard 0.5% glucose supply, Δ*pda1* mutants showed a consistent improvement in ethanol tolerance for both aerobic and anaerobic growth (*p*<0.005).

**Figure 6.**
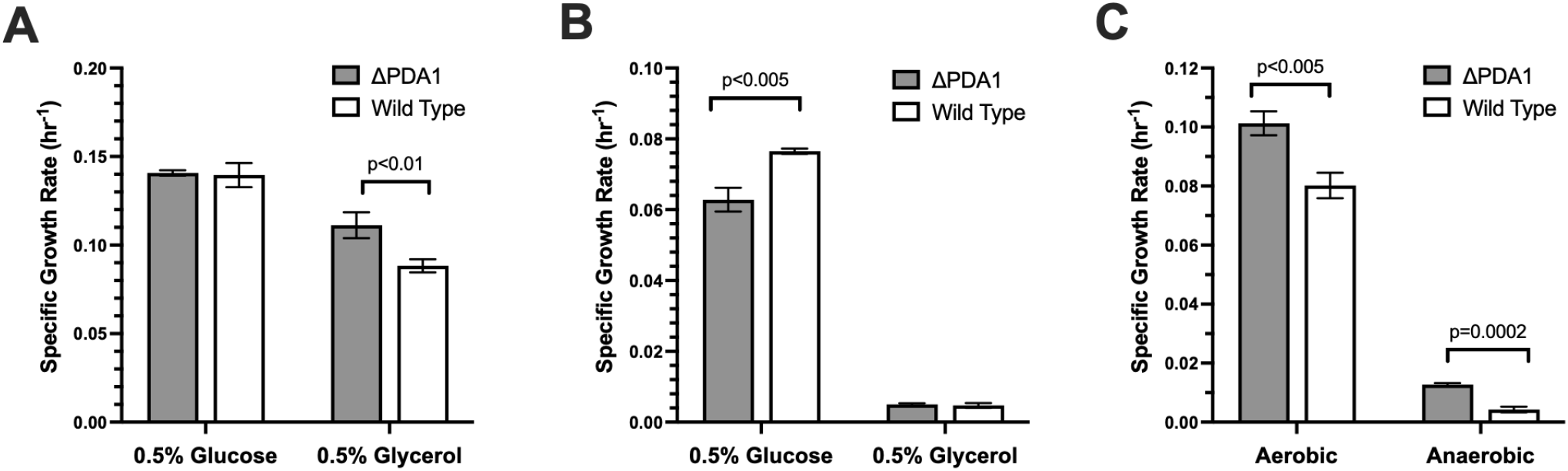
Glucose, glycerol, and ethanol metabolism among Δ*pda1* and wild-type yeast. **(A and B)** Mutant and wild-type yeast show differential preference for glucose and glycerol as sole carbon sources under aerobic (A) and anaerobic (B) environments. **(C)** Ethanol tolerance, measured by the growth rate of cultures grown on 0.5% glucose exposed to 5% added ethanol. Growth rates (hr^−1^) were calculated using OD_600_ measurements and averaged across 5 biological replicates (*n*=5). Uncertainty bars represent ±SD. Significance is determined by unpaired, two-tailed t-tests (\**p*<0.05, \*\**p*<0.01, \*\*\**p*<0.001).

Glycerol is generally considered to be a non-fermentable carbon source for most *S. cerevisiae* strains, meaning that oxygen is required for the metabolism of glycerol.^14^ Because a mutation in the *pda1* gene affects the aerobic metabolic pathway, it is surprising that a growth advantage is observed in Δ*pda1* mutant grown on glycerol. Because PDH is not directly involved in the conversion of glycerol into dihydroxyacetone phosphate (DHAP) to be used by the cell, it is hypothesized that the PDH deficiency in mutants alters gene expression patterns in other genes related to glycerol metabolism or that there was a change in flux of metabolites in the Δ*pda1* mutant. However, in both strains, glycerol was unable to be fermented anaerobically, demonstrating that such changes did not compensate for the inability of yeast to convert glycerol to ethanol (Fig. 6B).

Glycerol can be metabolized by wild-type cells directly into acetyl-CoA through PDH, and subsequently, enter the Krebs cycle and electron transport chain. Grown in glycerol, wild-type cells result in a greater net buildup of NADH compared to the Δ*pda1* mutant, due to the simultaneous procession of both the PDH and the PDH bypass pathways (Fig. 7A). The Δ*pda1* mutant degrades pyruvate through the PDH bypass, producing fewer molecules of NADH (Fig. 7B). Due to yeast’s inability to convert the excess NADH into NAD^+^ by fermentation, metabolism slows for wild-type cells when grown on glycerol as the sole carbon source.

**Figure 7.**
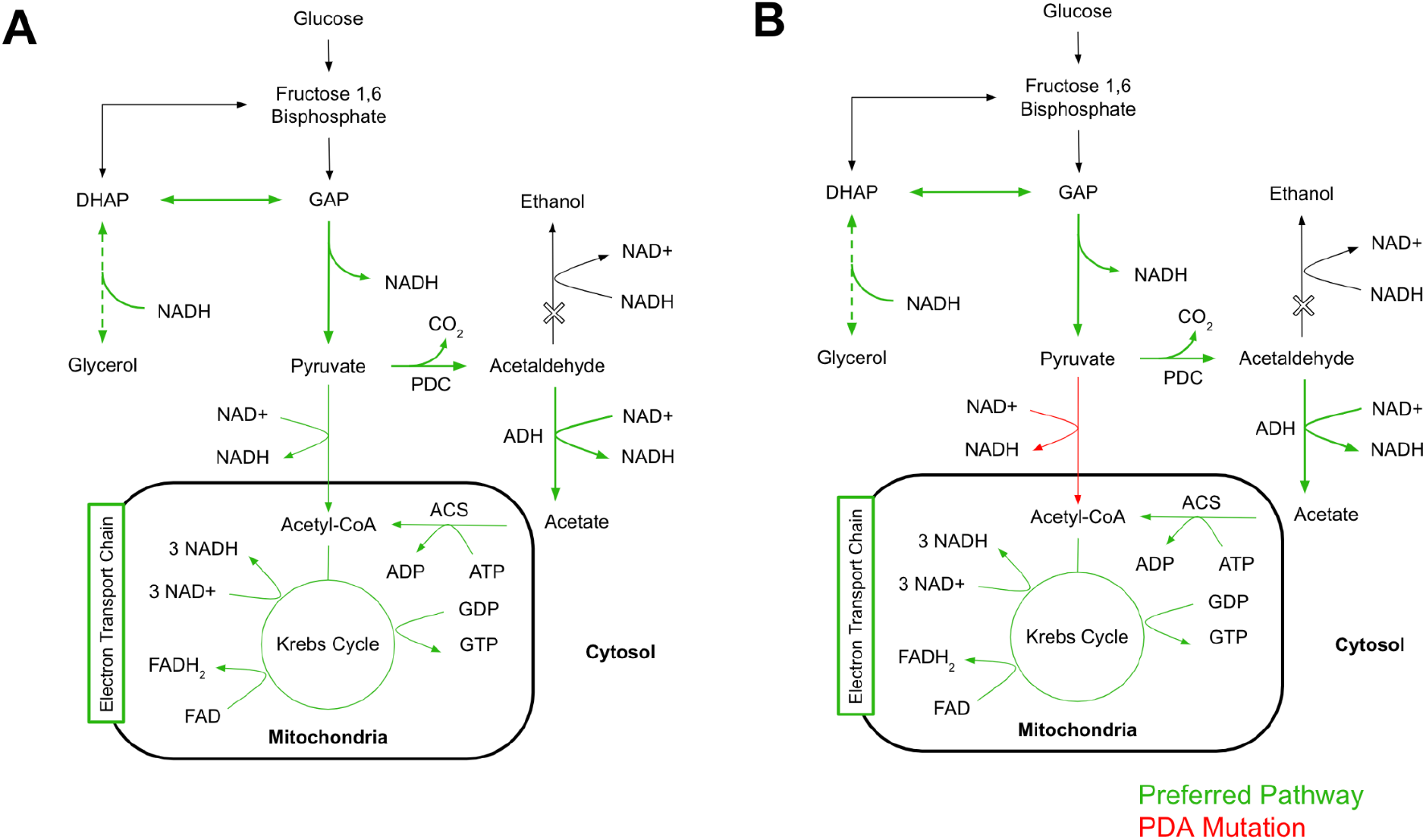
Proposed pathways of glycerol metabolism in wild-type and Δ*pda1* mutant yeast. Aerobic metabolism of glycerol was analyzed for the wild type (A) and mutant (B) grown in 0.5% glycerol as their sole carbon source. The conversion of pyruvate to acetyl-CoA is completed either through PDH or the PDH bypass in wild-type cells but only through the bypass in Δ*pda1* mutants. Ethanol is not produced as glycerol is a non-fermentable carbon source. Preferred pathways are denoted in green, and the product conversion rate is further distinguished by thickness. Red arrows denote a loss of function mutation of the PDH enzyme in Δ*pda1* mutants.^15^

### Implications of the Δpda1 Mutant Phenotype: A Novel Approach for Optimizing Bioethanol Synthesis

The main challenge hindering the optimization of *S. cerevisiae* phenotypes for the efficient production of bioethanol is the complexity of the genetic profile controlling yeast metabolism. Phenotypes are controlled by the concerted effects of hundreds of genes, and the targeting of any specific gene may not have a global effect that alters metabolic phenotypes substantively.^16^ Previous studies aiming to increase ethanol production in *S. cerevisiae* targeted genes primarily to suppress byproduct formation (biomass synthesis), or to increase ethanol tolerance. Both approaches were successful in enhancing ethanol production to an extent. However, a significantly favorable phenotypic change is often slightly counteracted by unfavorable side effects. For instance, a reduction in mutant growth rate in the absence of biomass limits the net ethanol production in yeast cultures despite an increase in fermentation rate.^6^ Similarly, attempts to increase cell viability by targeting ethanol tolerance genes successfully increased ethanol production by up to 25.7%, but enhancing tolerance alone was rarely sufficient to increase ethanol production past 25-30%.^17^ Global transcriptional rewiring through RNA polymerase II was able to increase ethanol production by up to 40% by simultaneously regulating multiple genes that contribute to ethanol tolerance.^16^

Consistent with the findings of previous studies, this study demonstrated that the observed increase in ethanol production is the product of two main phenotypic changes: the redirection of metabolic flux and an increase in ethanol tolerance. This study proposed Δ*pda1* as a candidate for the simultaneous optimization of both contributing phenotypes. Thus, the new Δ*pda1* mutant strain was able to increase ethanol production by 66.55% in 5% glucose, and up to two-fold in low-glucose cases (Fig. 3B). These results pointed to a new direction of addressing previous challenges, a promising advancement toward a more efficient synthesis of renewable biofuels.

## Conclusion

The deletion of *pda1* in *S. cerevisiae* BY4741 causes an absence of the pyruvate dehydrogenase enzyme, a key enzyme responsible for the conversion of pyruvate to acetyl-CoA at the entrance of the Krebs cycle. The PDH deficiency is shown to impede the aerobic growth of mutants, which can be attributed to the redirection of pyruvate to the less energetically favorable PDH bypass. Anaerobically, mutants supplement the lack of PDH and a nonfunctioning electron transport chain by using fermentation to maintain the NADH/NAD^+^ balance in the cell. Furthermore, the Δ*pda1* mutant cultures demonstrate an increased growth rate and a higher final OD under ethanol stress, as well as an improvement in glycerol metabolism. These combined changes in response to the PDH defect cause an observed increase in ethanol production ranging from 45.1% to 100.7% depending on the glucose supply. This increase in ethanol production is an important observation and will aid future sustainable production of renewable biofuels from glucose and other carbon sources. These data also provide context for future research in creating strains beneficial for ethanol production. Further studies can be conducted to determine the cause of the ethanol tolerance increase by mapping the expression levels of genes proposed to play a role in complementing the PDH deficiency in mutants, such as the *PDC, ADH*, and *ACS* genes involved in the PDH bypass.

## Acknowledgments

The authors would like to thank the Bose Lab of Washington University in St. Louis for providing all the needed equipment.

